# Molecular Evidence of Internal Carbon-Driven Partial Denitrification Annamox (PdNA) in a mainstream Pilot A-B System Coupled with Side-stream EBPR treating municipal wastewater

**DOI:** 10.1101/2023.03.28.534645

**Authors:** Da Kang, IL Han, Jangho Lee, Kester McCullough, Guangyu Li, Dongqi Wang, Stephanie Klaus, Ping Zheng, Varun Srinivasan, Charles Bott, April Z. Gu

## Abstract

Achieving mainstream short-cut nitrogen removal via nitrite has been a carbon and energy-efficient goal which wastewater engineers are dedicated to explore. This study applied a novel pilot-scale A-B-S2EBPR system process integrated with sidestream enhanced biological phosphorus removal) to achieve the nitrite accumulation and downstream anammox for treating municipal wastewater. Nitrite accumulated to 5.5 ± 0.3 mg N/L in the intermittently aerated tanks of B-stage with the nitrite accumulation ratio (NAR) of 79.1 ± 6.5%. The final effluent concentration and removal efficiency of total inorganic nitrogen (TIN) were 4.6 ± 1.8 mg N/L and 84.9 ± 5.6%, respectively. Batch nitrification/denitrification activity tests and functional gene abundance of ammonium oxidizing bacteria (AOB) and nitrite oxidizing bacteria (NOB) suggested that the nitrite accumulation was mostly caused by partial denitrification without NOB- selection. The unique features of S2EBPR (longer anaerobic HRT/SRT, lower ORPs, high and more complex VFAs etc.) seemed to impact the nitrogen microbial communities: the conventional AOB kept at a very low level of 0.13 ± 0.13% during the operation period, and the dominant candidate internal carbon-accumulating heterotrophic genera of *Acinetobacter* (17.8 ± 15.5)% and *Comamonadaceae* (6.7 ± 3.4%) were highly enriched. Furthermore, the single-cell Raman spectroscopy-based intracellular polymer analysis revealed the dominate microorganisms that could utilize polyhydroxyalkanoates (PHA) as the potential internal carbon source to drive partial denitrification. This study provides insights and a new direction for implementing the mainstream PdNA short-cut nitrogen removal via incorporating S2EBPR into sustainable A-B process.

## 1. Introduction

The conventional activated sludge processes are energy-intensive, consuming as much as 25-40% of the total operational costs in a typical wastewater treatment plant (WWTP) (Ehrhard and Murphy, 2009) and 3% of the generated electricity in the United States (Logan and Elimelech, 2012; McCarty et al., 2011). To transit WWTPs towards energy-neutral and resource recovery-oriented plants, the A-B processes have been developed where the A-stage is designed for carbon capture for energy recovery and B-stage is for nutrient removal and recovery (Liu et al., 2019; Rahman et al., 2020).

Since most influent carbon would be firstly captured at A-stage, B-stage is preferable to implement the short-cut nitrogen removal processes via nitrite instead of nitrate. As successfully demonstrated in treating ammonium-rich streams (e.g., sludge digestate, leachate and industrial wastewater), the partial nitritation/anammox (PN/A) processes could save 100% carbon, 60% aeration and reduce 80% sludge production (Kartal et al., 2010; Lackner et al., 2014; Van der Star et al., 2007). However, the realization of short-cut nitrogen removal in mainstream B-stage is still the main challenging aspect or sometimes bottleneck for municipal low-strength wastewaters, where the sustainable supply of the intermediate nitrite could not be easily acquirable by controlling temperature, free ammonia (FA), free nitrous acid (FNA), dissolved oxygen (DO), etc. to suppress the nitrite oxidizing bacteria (NOB) over ammonium oxidizing bacteria (AOB) (Cao et al., 2017; Laureni et al., 2016).

Recently, partial denitrification (PDN) was proposed and demonstrated as a practically more promising and stable alternative to achieve the mainstream nitrite shunt for the low-strength municipal wastewaters (Du et al., 2015; Ma et al., 2017). The integration of PDN/anammox could theoretically reduce 47.7% oxygen demand, 54.2% COD requirement and 66.3% sludge production compared to the conventional nitrification/denitrification process (Zhang et al., 2019). To date, the type of carbon sources, limited ratio of carbon to nitrogen (C/N), high pH, high salinity, and the alternating feast-famine regime were reported to be the selection pressure to achieve successful enrichment of partial denitrifying microorganisms (Zhang et al., 2020). Generally, acetate or the sludge fermentation liquid was proved to be the most ideal carbon source to achieve the efficient PDN process in the lab-scale reactors than any other type (glucose, methanol, ethanol, etc.) (Cao et al., 2013; Le et al., 2019; Li et al., 2018a; Zhang et al., 2020). The genera of *Thauera* and *Halomonas* were respectively reported to be highly enriched in acetate-fed denitratation system by different research groups via amplicon sequencing and suspected to be the progressive onset denitrifiers with much higher functional gene abundance and transcription of nitrate reductase than nitrite reductase (Li et al., 2016; Qian et al., 2019). In addition, the endogenous partial denitrification was also found driven by microorganisms capable of transforming the external carbon source to the internal carbon storage. In a *Halomonas-*dominated denitratating culture, with the sufficient acetate supply (C/N=5), it was observed that the poly-β-hydroxy-butyrate (PHB) accumulated first during the nitrate to nitrite phase, and then consumed with the reduction of nitrite (Li et al., 2016). However, in another studies, acetate was firstly taken up accompanied with the glycogen consumption and polyhydroxyalkanoates (PHA) accumulation during the anaerobic phase in a lab-scale reactor, and then utilized to reduce nitrate to nitrite via bulk chemical analysis, resulting the enrichment of denitrifying glycogen accumulating organisms (DGAOs) (Ji et al., 2018; Zeng et al., 2003b). Even though, most of the PDN study was still at the early stage in lab-scale and the demonstration of PDN process at pilot- or full-scale was only be reported recently by a few studies (Fofana et al., 2022; Zhao et al., 2021). Pure culture studies of denitrifying bacteria implied that nitrate reductase has a more competitive advantage for electrons over nitrite reductase leading to the transient accumulation of extracellular nitrite during denitrification (Almeida et al., 1995; Glass and Silverstein, 1998), this provide the fundamental basis to enhance the nitrite accumulation by either sub-sufficient carbon supply or lower electron donor flux rates by internal PHA hydrolysis to favor the PDN process (Zeng et al., 2003a). Particularly, the identities and roles of specific organism that are capable of utilizing internal carbon (i.e., PHA) storage to facilitate the PDN process have hardly been investigated.

The emerging alternative side-stream enhanced biological phosphorus removal (S2EBPR) processes utilizes the hydrolysis and partial fermentation of return activated sludge (RAS) to generate internal carbon, and provides influent carbon-independent, therefore more controlled, stable and favorable phosphorus accumulating organisms (PAOs) enrichment conditions to enable and enhance the EBPR performance and stability (Barnard et al., 2017; Gu et al., 2018; Onnis-Hayden et al., 2020; Srinivasan et al., 2021; Wang et al., 2019a). Another unique observation is that S2EBPR enriched for higher abundance of PHA-accumulating organisms than conventional EBPR systems (Wang et al., 2021). Therefore, we hypothesized that the incorporation of S2EBPR may potentially facilitate mainstream PDN by enriching for higher abundance of PHA-accumulating organisms than conventional BNR processes, which would favor internal carbon-driven PDN. Our team have recently reported the feasibility of combining the S2EBPR with A-B stage to achieve short-cut nitrogen removal and phosphorus removal (Stephanie et al., 2018). In this study, we investigated the nitrite accumulation predominantly attributed to PDN rather than PN (NOB out-selection was not observed) in this pilot A-B-S2EBPR system treating municipal wastewater treatment. Process performance, carbon and nitrogen mass balance, batch activity tests were performed to investigate the nature and cause of the nitrite accumulation. 16S rRNA gene amplicon sequencing and qPCR were applied to detect the microbial communities and phylogenetic abundance of conventional nitrifiers (AOB and NOB). In addition, single-cell Raman spectroscopy (SCRS) and hierarchical clustering analysis were conducted to obtain the phenotypic profiling of the dominant bacteria and quantify the intracellular carbon polymers (including PHA, glycogen, polyphosphate, etc.) dynamics within the dominant operational phenotypic units (OPUs) during the partial denitrification process. This research advances our understanding and provides insights towards a new direction for integrating the S2EBPR to the sustainable A-B process to realize the mainstream short-cut nitrogen removal.

## 2. Materials and methods

### 2.1. Pilot A-B-S2EBPR system

The pilot plant operated by Hampton Roads Sanitation District (HRSD) is located at the Chesapeake-Elizabeth WWTP in Virginia Beach, VA, USA. The plant runs in an A-B process consisting of the high-rate activated sludge (HRAS) process at A-stage followed by B-stage designing for achieving nitrite accumulation, and then combined with the anammox moving bed biofilm reactor (MBBR) polishing step to accomplish short-cut nitrogen removal (Fig. 1). The detailed parameters of A-B process have been introduced elsewhere (Klaus et al., 2020; Printz, 2019). Briefly, the B-stage includes an anaerobic selector with a volume of 53 L, and followed by four identical continuous stirred tank reactors (CSTRs) in series with the total volume of 600 L. The CSTRs are intermittently aerated via Ammonia versus NO_x_ (nitrite+nitrate) (AvN) control strategy (Regmi et al., 2014) with hydraulic retention time (HRT) of 5 h and sludge retention time (SRT) of 8.3 ± 1.3 days. The pilot plant also includes a 340 L MBBR loaded with K3 anammox biofilm carriers (AnoxKaldnes, Sweden) and a 174 L side-stream biological phosphorus remover (SBPR) receiving part of return activated sludge (RAS) and fermentate from A-stage wasted activated sludge (WAS) fermenter to enhance the phosphorus removal. For this study, the pilot-plant was operated for 100 days and the detailed operational and performance parameters were summarized in Table S1.

**Fig. 1.**
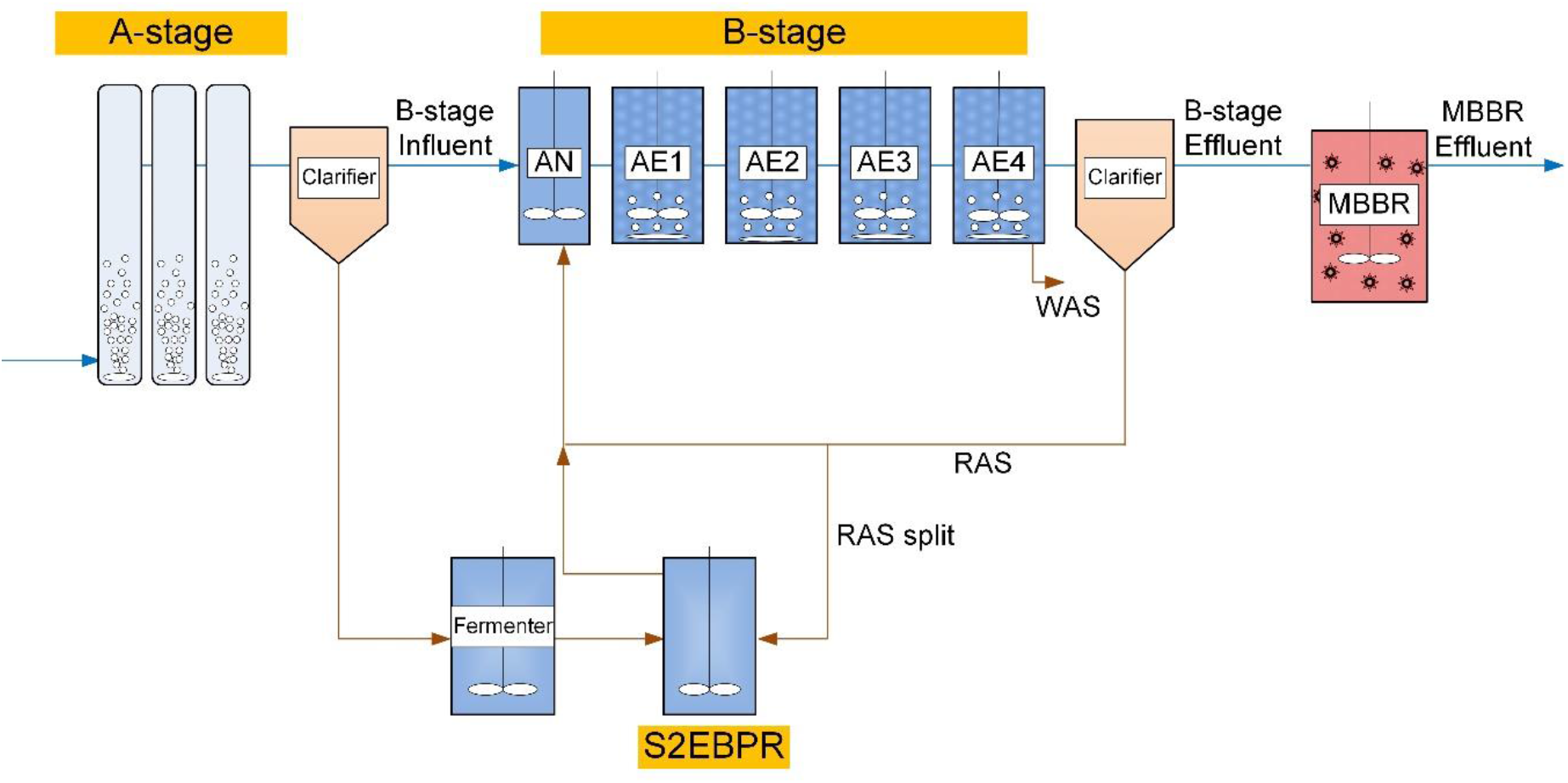
Schematic configurations of the pilot A-B-S2EBPR system adapted from (Klaus et al., 2020) (RAS-return activated sludge; WAS-wasted activated sludge; AN- anaerobic selector; AE-aerobic continuous stirred tank reactors; S2EBPR-sidestream enhanced biological phosphorus remover; MBBR-moving bed biofilm reactor).

### 2.2. Batch nitrification activity tests

The batch nitrification activity tests were performed weekly on site. About 4 L mixed liquor suspended solids (MLSS) were withdrawn from the 4^th^ aerobic tank at B-stage and pre-aerated for 30 minutes to oxidize the residue carbon, and then spiked 25 mg N/L ammonium chloride, pH was maintained around 7, and DO was around 3-5 mg/L. The samples were taken every 15 minutes for a total of 1 h, filtered by 0.45 μm filter and then analyzed for NH_4_^+^-N, NO_2_^-^-N, NO_3_^-^-N concentrations by colorimetric method using HACH TNT kits (HACH Loveland, CO). At the end of batch tests, the total suspended solids (TSS) was measured according to the standard methods (APHA, 2005). AOB activity was calculated as the sum of nitrite and nitrate production rates and NOB activity was calculated as the nitrate production rates by linear regression.

### 2.3. Batch denitrification activity tests

The batch denitrification activity tests were performed weekly on site. About 2 L RAS from B-stage secondary clarifier was mixed with A-stage effluent (soluble COD < 100 mg/L) and 20 mg N/L nitrate was spiked to perform the denitrification test. The samples were taken every 15 minutes for a total of 1 h, filtered by 0.45 μm filter, and then analyzed for COD, NH_4_^+^-N, NO_2_^-^-N, NO_3_^-^-N and TSS concentrations using the same methods as previously described. The nitrate reduction and nitrite accumulation rates were respective calculated by linear regression.

In addition to the routine denitrification batch activity tests, the specific denitrification activity test with pre-incubation by fermentate was performed on site on day 63 to maximize the accumulation of internal carbon source (like PHA) and identify which promoting the denitrification. Sludge from the anaerobic selector was incubated with A-stage WAS fermentate (VFA = 950 ± 40 mg COD/L) overnight, and then washed twice with B-stage secondary effluent (COD = 83.7 ± 11.5 mg/L). 10 mg N/L nitrate was spiked at first and the water and sludge samples were taken every 30 minutes for a total of 2 h. The analytical methods for soluble COD (sCOD), NH_4_^+^-N, NO_2_^-^-N, NO_3_^-^- N and TSS concentrations were the same as previously described. PHA including poly- β-hydroxy-butyrate (PHB), poly-β-hydroxy-valerate (PHV), poly-β-hydroxy-2- methyl-valerate (PH2MV) were extracted with a 3 h digestion time and 3% sulfuric acid, and analyzed by gas chromatography-mass spectrometry (GC-MS) (Lanham et al., 2013). The glycogen was extracted with a 2 h digestion time and 0.9 M hydrochloric acid, and analyzed by liquid chromatography-tandem mass spectrometry (LC-MS/MS) (Wang et al., 2019a).

### 2.4. DNA extraction and 16S rRNA gene amplicon sequencing

Genomic DNA of sludge samples from the B-stage during the operation were extracted using the DNeasy PowerSoil Pro Kit (Qiagen, USA). The V4 region of the 16S rRNA gene was amplified using the primers 515F/806R and then the amplicons were sequenced on the Illumina’s MiSeq with V2 chemistry using paired-end (2 x 250) sequencing. The raw paired-end reads were assembled for each sample and analyzed following the standard operating procedure by Mothur software (Kozich et al., 2013). High-quality reads were obtained after quality control and chimera screening and then clustered at a 97% similarity to obtain the operational taxonomic units (OTUs). The representative sequences of each OTU were annotated based on Silva database (Quast et al., 2012). The raw reads were deposited into the Sequence Read Archive (SRA) database in National Center for Biotechnology Information (NCBI) with the accession number of PRJNA866204.

### 2.5. Quantitative polymerase chain reaction (qPCR)

The gene abundance of total bacteria and nitrifiers were determined by the quantitative polymerase chain reaction (qPCR) using the CFX real-time PCR detection system (Bio-Rad, USA). The primer sets chosen for AOB, NOB, complete ammonia oxidation (Comammox) and total bacteria were amoA-1F/2R (Rotthauwe et al., 1997), nxrB-169F/638R (Pester et al., 2014), amoB 148F/485R (Cotto et al., 2020) and 16S rRNA-341F/534R (He et al., 2007), respectively. The sequence and thermocycling conditions for each primer were in supplementary information (Table S2). The qPCR reaction mixture in total of 20 μL containing 1 μL DNA template, 1 μL each primer, 7 μL RNase-free water and 10 μL iQ SYBR^○R^ Green Supermix (Bio-Rad, USA). The standard curves were constructed from a series of 10-fold dilutions of a plasmid DNA obtained by TOPO TA cloning (ThermoFisher, USA). The R^2^ of standard curves were above 0.99 and the amplification efficiency could reach around 95%.

### 2.6. Single-cell Raman spectroscopy (SCRS)

The sludge samples at time 0, 0.5 h, 1 h, 2 h during the pre-incubated denitrification batch test with fermentate were respectively collected to acquire single-cell Raman spectra for phenotypic profiling of the microbial community and the details of Raman phenotyping analysis can be referred to the previous reference (Li et al., 2018b; Majed and Gu, 2010). Briefly, the sludge samples were washed, diluted, homogenized by a 26-gauge syringe needle and dried on optically polished CaF_2_ circular windows (Crystran, UK). Raman spectra of single-cell were scanned from 400 to 1800 cm^-1^ by a LabRam HR Evolution Raman microscope equipped with a magnification of x50 objective (HORIBA, Japan). The laser wavelength was 532 nm and the gratings were 600 grooves/mm with an acquisition time of 20s per spectrum. The spectra data were processed with the smoothing and filtering, background subtraction, baseline correction steps by the software LabSpec 6 (Detailed key parameters can be found in supporting information Text S1). The statistical sufficiency of the sampling size was recently evaluated by kernel divergence computing method which showed that an approximated sampling size of 50 or 100 spectra for full-scale EBPR systems at 5% or 2% operational phenotypic units (OPUs) cluster resolution (Li et al., 2022; Majed et al., 2012). A total of 131, 190, 197 and 209 Raman spectra of single cells were obtained finally at each sampling time. The presence of PHA in single cells was identified by the signature peaks in the range of 1715-1740 cm^-1^ based on previous study (Majed and Gu, 2010). The relative intensity of PHA for single cell was normalized by the intensity of amide I vibration at 1640-1690 cm^-1^ as the biomass marker. The hierarchical clustering analysis (HCA) was applied on all of the single-cell Raman spectra from activated sludge samples to obtain phenotypic profiles based on OPUs, and the cosine similarity (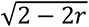, r-correlation efficient) was used to measure metrics between two samples, average linkage was applied to quantify dissimilarities between two clusters and the cutoff threshold for OPUs was set at 0.82 according to previous experiments (Li et al., 2018b).

## 3. Results and discussion

### 3.1. Nitrite accumulation in A-B-S2EBPR system

The A-B-S2EBPR pilot plant was operated for around 100 days for this period of the study. The average ammonium concentration of B-stage influent was 30.9 ± 3.7 mg N/L and the effluent ammonium was 6.5 ± 1.4 mg N/L (Fig. 2A). Nitrite accumulation could be detected in B-stage effluent and reached the highest around 5.5 ± 0.3 mg N/L (Fig. 2B), meanwhile, the B-stage effluent nitrate decreased gradually from 8.1 mg N/L to 1.0 mg N/L (Fig. 2C). The nitrite accumulation ratio (NAR) calculated by the ratio of nitrite to the sum of nitrite and nitrate of B-stage effluent could reach 79.1 ± 6.5%. With the further polishing step by MBBR, the overall effluent concentration of total inorganic nitrogen (TIN) was 4.6 ± 1.8 mg N/L with removal efficiency of 84.9 ± 5.6% (Fig. 2D).

**Fig. 2.**
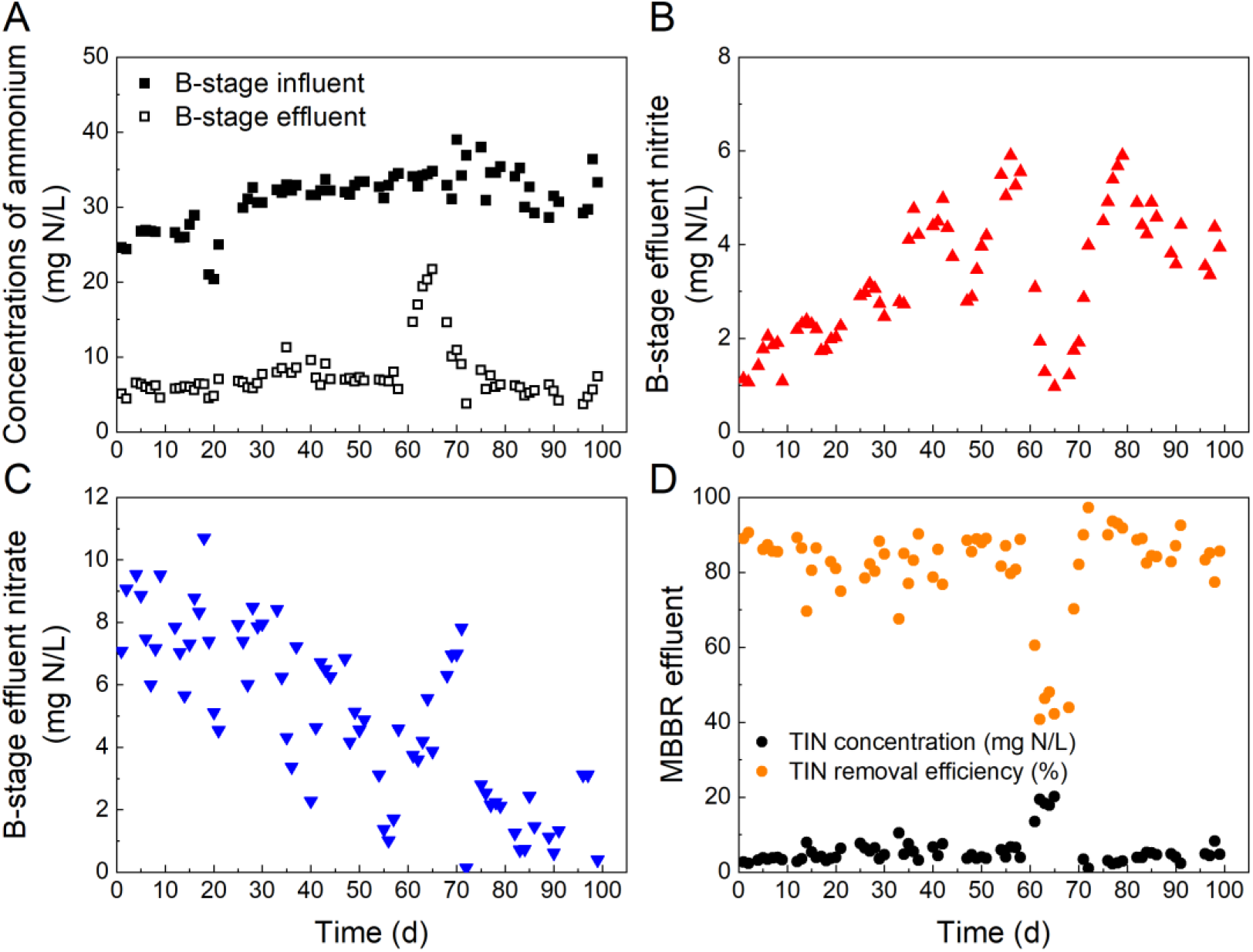
Nitrogen removal performance of A-B-S2EBPR system (A-ammonium concentrations of B-stage influent and effluent; B-nitrite concentration of B-stage effluent; C- nitrate concentration of B-stage effluent; D-total inorganic nitrogen (TIN) concentration and removal efficiency from B-stage influent to MBBR (Anammox reactor) effluent; the sampling positions of B-stage influent, B-stage effluent, MBBR effluent were marked in Fig. 1; the performance collapse between day 60 and day 70 was caused by operational issues).

Carbon and nitrogen mass balance were further calculated for the A-B-S2EBPR pilot system during the operation period where the nitrite accumulation occurred. For soluble carbon (sCOD), the integration of A-stage WAS fermenter largely supplement the carbon input to the S2EBPR and then to the B-stage tanks, enhanced from solely B-stage influent of 343.2 g/d to the total of 569.3 g/d (Fig. 3A). For inorganic nitrogen, B-stage influent was the main source of nitrogen input to the down-steam, accounting for 87.5% of the total nitrogen input. But it was noted that most carbon (59.4%) from fermentate was captured in S2EBPR and the residuals went into the anoxic and aerobic tanks of B-stage were very limited with the average soluble COD concentration of only 58.3 ± 13.3 mg/L, leading to the low TIN removal efficiency of 25.4%. However, the nitrite mass flow gradually increased along the B-stage tanks from 5.4 g/d to 28.5 g/d and the NAR could reach 41.7% at the 4th AE tank, which could provide the intermediate nitrite to enable the short-cut nitrogen removal of the subsequent MBBR. MBBR played the main function of TIN removal with the removal efficiency of 69.8% (Fig. 3B).

**Fig. 3.**
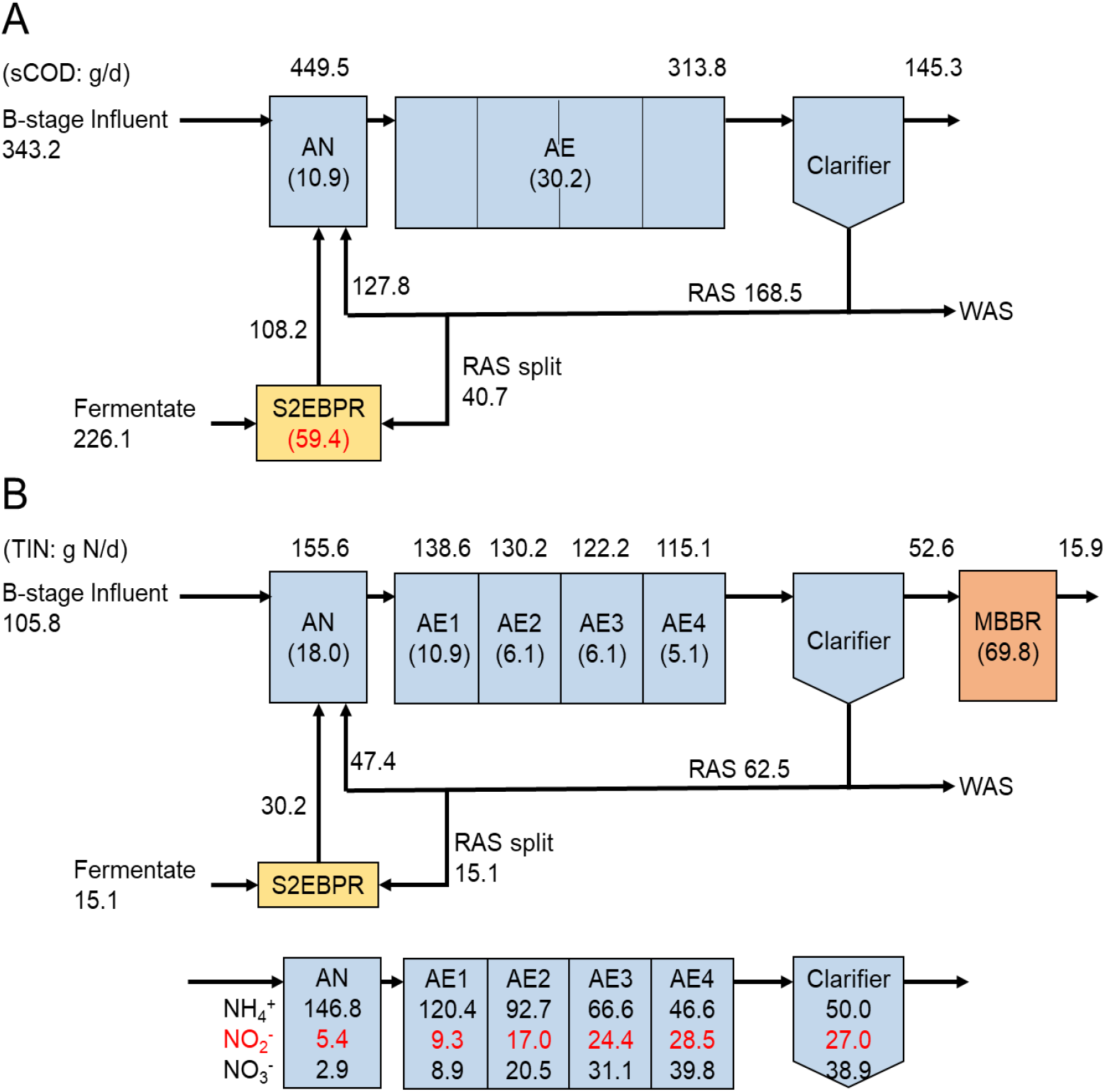
Carbon (A) and nitrogen (B) mass balance of A-B-S2EBPR system (all units are in g/d; values in brackets represent the percentage (%) of nitrogen conversion in the operational unit).

### 3.2. Nitrite accumulation was mainly caused by partial denitrification

To investigate the cause of nitrite accumulation in the B-stage, we simultaneously evaluated multiple aspects including AOB/NOB activities and denitrification activities, *amo* and *nxr* gene abundance indicative of AOB and NOB, as well as nitrogen species analysis during the intermittent aeration cycles in the B-stage reactor (Fig. 4).

**Fig. 4.**
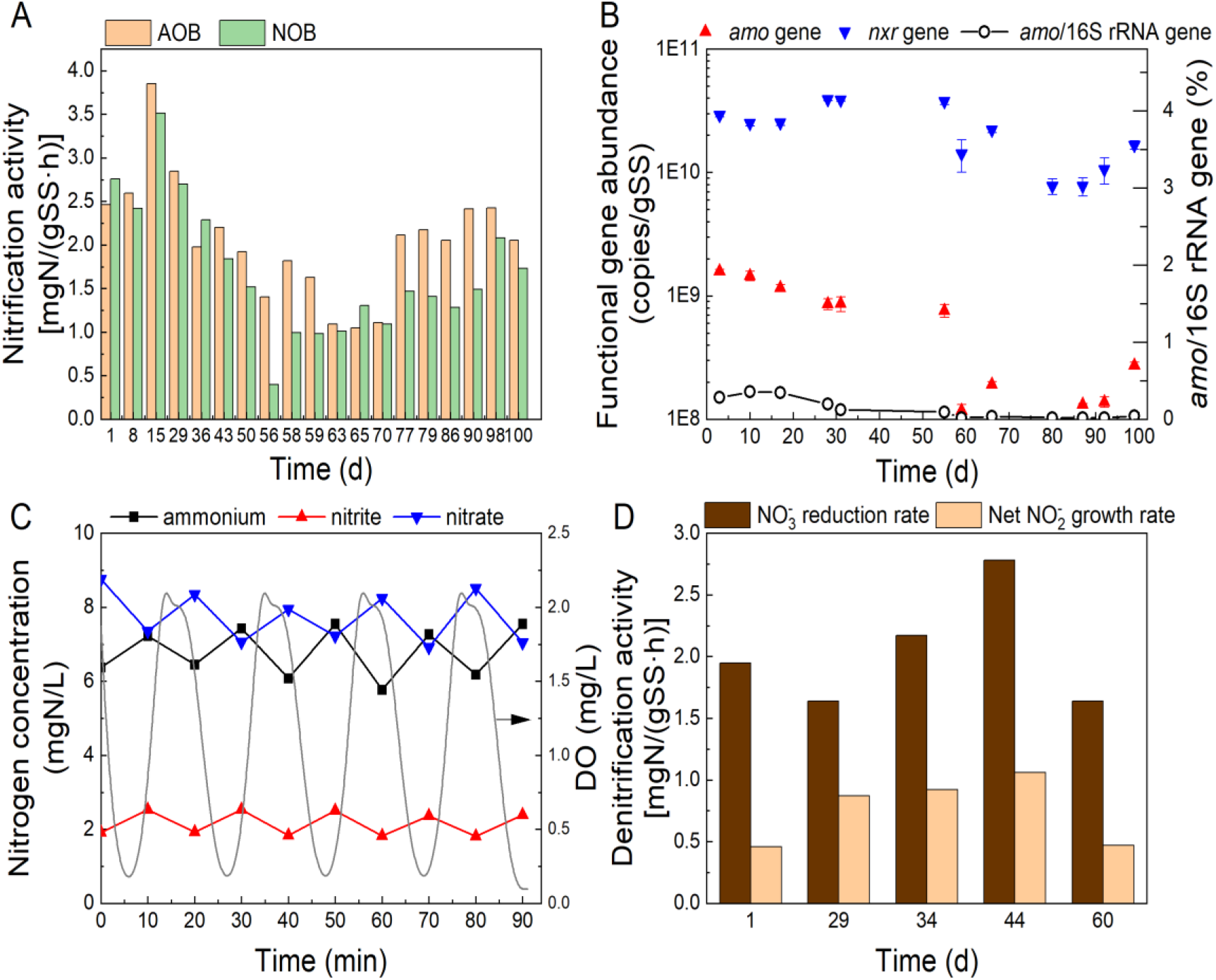
Temporal nitrification and denitrification activities, abundance of nitrogen microorganisms and nitrogen species transformation pattern during the study supporting the PDN process. (A) Nitrification activities; (B) Functional gene abundance of AOB and NOB; (C) Typical time cycle of nitrogen species and DO variation in the intermittently aerated 4th AE tank of B-stage and (D) Denitrification activities.

The nitrification batch activity tests showed that no significant difference could be detected between the specific activity of AOB and NOB during the operation period where nitrite accumulated (n = 16, *p* = 0.23, one-way ANOVA) (Fig. 4A). Moreover, the functional gene abundance of AOB (*amo* gene) and NOB (*nxr* gene) were detected by qPCR and the results showed that the average abundance of nitrate-reducing *nxr* gene [(2.3 ± 1.2) × 10^10^ gene copies/g SS] was two magnitudes higher than that of ammonia-oxidizing *amo* gene [(6.3 ± 5.5) × 10^8^ gene copies/g SS] (Fig. 4B), which indicated that NOB-out-selection was not achieved in this system. It was also noted that the relative abundance of AOB (calculated as the ratio of *amo* gene copies to 16S rRNA gene) was as low as 0.13 ± 0.13%. The relative abundance of comammox was also at very low level of 0.12 ± 0.11% (Fig. S1). These evidenced that PN was unlikely the main process that led to nitrite accumulation. The denitrification batch activity tests showed that with the reduction of nitrate, the nitrite gradually accumulated with the net growth rate of 0.68 ± 0.34 mg N/(g SS·h) (Fig. 4D), which was consistent with the periodic time cycle of nitrogen conversion in the B-stage AE tank. The nitrite accumulation only occurred in the anoxic phase with concurrent nitrate decline and nitrite was oxidized in the aerobic phase (Fig. 4C). These collective evidences led to our conclusion that partial denitrification was the main pathway leading to the observed nitrite accumulation.

### 3.3. Partial denitrification associated with internal carbon utilization

Two observations that led to our hypothesis that PHA could be the potential internal carbon to drive the partial denitrification in the A-B-S2EBPR pilot system. First, according to the mass balance analysis in our pilot system, most of carbon from fermentate was captured in A-stage and then fed into S2EBPR and very low soluble COD residual entering into the anoxic and aerobic tanks of B-stage with average concentration of only 58.3 ± 13.3 mg/L (Fig. 2). Second, our previous study via Raman-based observation suggested that the PHA-containing cells were significantly higher in the S2EBPR system in comparison to those in the conventional EBPR systems (Wang et al., 2021; Wang et al., 2018). Coupling of internal carbon (PHA) utilization and partial denitrification were also observed by others (Ji et al., 2017; Krasnits et al., 2013; Tu et al., 2019; Wang et al., 2019b). To test this hypothesis, we performed denitrification batch tests with no external carbon addition, where sludge samples that were pre-incubated with fermentate to enrich for PHA-accumulating organisms and then washed prior to the batch test to remove all residue soluble carbon. The results showed that without adding external carbon (the bulk residual sCOD fluctuated between 45-80 mg/L), nitrite accumulated from 1.4 mg N/L to 2.8 mg N/L with nitrate reduction from 9.4 mg N/L to 0.8 mg N/L (Fig. 5). In contrast, in a parallel denitrification test, no nitrite accumulation was observed when external carbon acetate was added, and PHA accumulated instead of being consumed (Fig. S2). Bulk measured PHA declined from 54.6 to 44.8 mg/g TS concurrently with nitrate decline, while the concentration of glycogen fluctuated between 65.0 to 88.7 mg/g TS. The stoichiometric analysis confirmed that the ratio of PHA consumption to nitrate reduction was 4.17 g/g N, which was comparable with the theoretical PHB-driven denitrification ratio of 1.71-4.39 g/g N (Hiraishi and Khan, 2003).

**Fig. 5.**
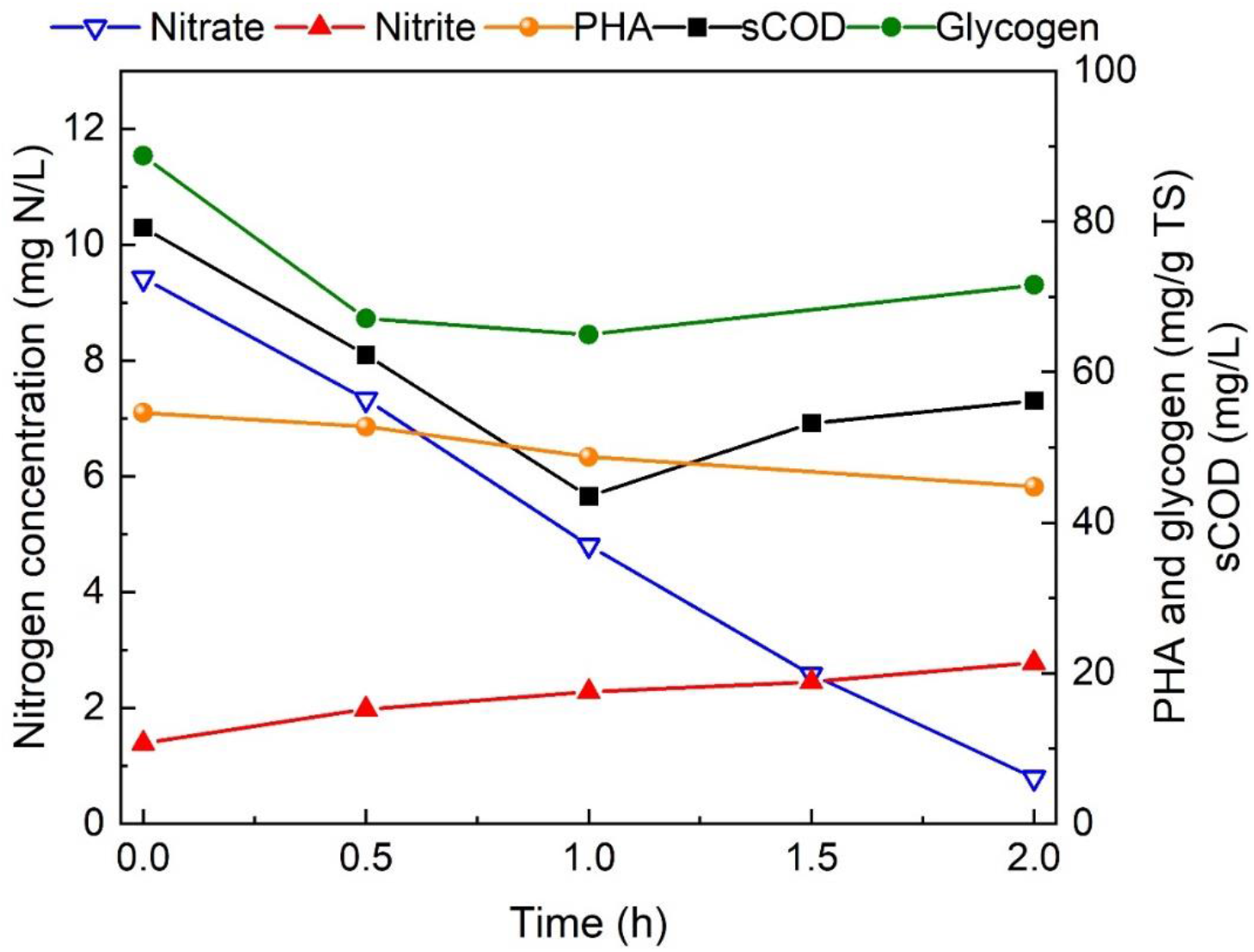
Variations of nitrogen and intracellular compounds during fermentate pre-incubated denitrification batch activity test.

### 3.4. Potential roles of PHA-accumulating microorganisms in PDN in the pilot A-B-S2EBPR system

The microbial communities of the A-B-S2EBPR pilot system were investigated by temporal 16S rRNA gene amplicon sequencing. At the phylum level (Fig. 6A), *Proteobacteria* was the most dominated with the relative abundance of 51.4%, where most denitrifiers belong to the different classes of *Proteobacteria* (α-, β-, γ-, and ε- *Proteobacteria*) (Braker and Conrad, 2011). *Firmicutes*, as the well-known fermenters, were the second abundant phyla (23.4%) and often widely distributed in anaerobic digesters (Garcia et al., 2011; Yang et al., 2014). At genus level (Fig. 6B), *Acinetobacter* was the most abundant genera with the relative abundance of (17.8 ± 15.5)%, followed by *Comamonadaceae* (6.7 ± 3.4%), *Beijerinckiaceae* (5.3 ± 4.5%) and *Enterobacteriaceae* (5.1 ± 2.9%). None of the reads belonging to the conventional AOB were sequenced and *Nitrospira* was the only detectable NOB with the percentage as low as 0.2 ± 0.2%. This was consistent with the activities and gene abundance results of AOB and NOB discussed previously (Fig. 4A & 4B). It is also conceivable that there exist other unknown AOB that was not captured by the sequencing analysis that performed ammonia oxidation. The surprisingly high abundance of *Acinetobacter* species suggested that it may play the functional role in our system. A number of *Acinetobacter* species were reported to have the ability of simultaneous heterotrophic nitrification and aerobic denitrification (Chen et al., 2019; Huang et al., 2013; Su et al., 2015; Yang et al., 2019; Yao et al., 2013), which may provide one possible explanation for (at least partially) why the relatively high ammonium oxidation activities observed (Fig. 4A) when almost no known autotrophic nitrifiers was detected. The pure culture of *Acinetobacter junii* isolated from activated sludge was proved to not only oxidize ammonium, but can also reduce nitrate with nitrite accumulation observed (Ren et al., 2014). Furthermore, *Acinetobacter* sp. were demonstrated to have the ability of synthesizing PHA as the internal carbon storage (Anburajan et al., 2019; Yang et al., 2013). *Comamonadaceae* are a large and diverse bacterial family of the *Betaproteobacteria* and were affiliated with denitrifying groups, especially a number of PHA-degrading denitrifying bacteria have been assigned phylogenetically to the family *Comamonadaceae* (Hiraishi and Khan, 2003). *Comamonadaceae* were also detected to be dominant in the partial denitrification system (Du et al., 2016; Wu et al., 2018). In addition, the family *Beijerinckiaceae* include a metabolically diverse aerobic bacteria from obligate methanotrophs to chemoorganoheterotrophs and have a genera trait of forming poly-b-hydroxybutyrate (PHB) granules (Marín and Arahal, 2014). The family *Enterobacteriaceae* are facultative anaerobes with the ability of fermenting glucose to produce lactic acid and reducing nitrate to nitrite (Brenner and Farmer III, 2015).

**Fig. 6.**
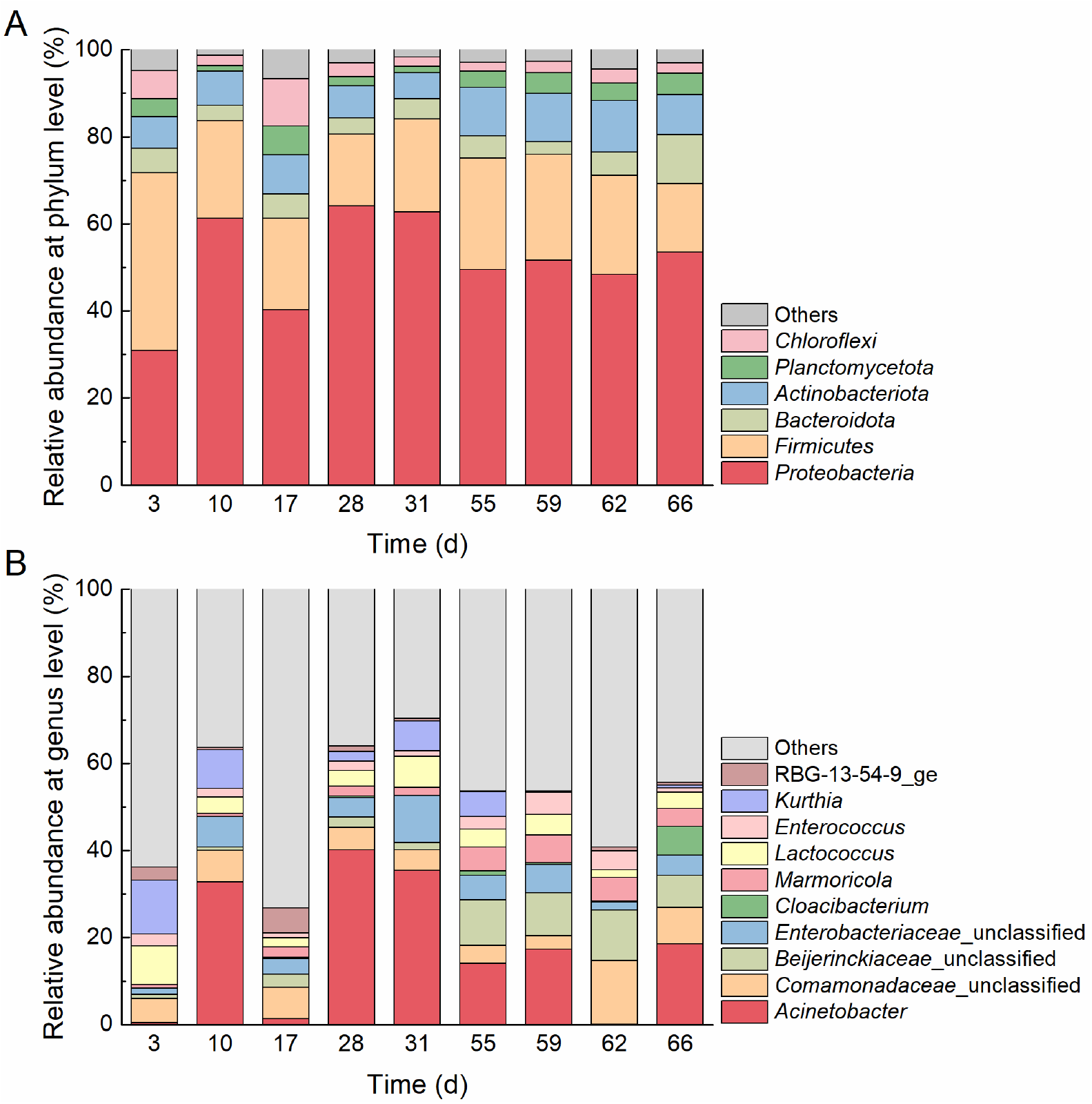
The relative abundance of microbial communities during the operation period at phylum (A) and genus (B) level.

To further investigate if PHA likely acted as the internal carbon source for denitrification, the sludges at different time of denitrification batch test pre incubated by fermentate were sampled and detected by single-cell Raman spectroscopy (SCRS) to identify the main operational phenotypic units (OPUs) in the biomass via HCA analysis. Previous studies indicated that single-cell Raman signature allowed for identification of organism at resolution level comparable to species or even strain level (Hutsebaut et al., 2006; Li et al., 2018b). With the aim to obtain further insights in the role of PHA and PHA-accumulating organisms in partial denitrification, single-cell Raman spectroscopy (SCRS) was applied to phenotyping the microbial community and to reveal the variations of relevant intracellular storage polymers during the batch denitrification test (Li et al., 2018b; Majed et al., 2012). Three main dominant operational phenotypic units (OPUs, named OPU 1-3) were detected by hierarchical clustering analysis (HCA) of SCRS spectrum (Fig. 7A). OPU1 (operational phenotypic unit) was the most dominant one with the relative abundance of 5.26-25.89% based on the phenotypic analysis (Fig. 7B), and the intracellular PHA were only detected in cells within the OPU1. The average relative Raman intensity of PHA in the cells of OPU1 declined from 2.51 to 1.54 (Fig. 7C). To get insights on the phylogenetic identities of cells in OPU1, we clustered the single cell Raman spectra obtained from the biomass in our system with available spectrum obtained from pure culture of *Acinetobacter junii* and *Comamonas testosteroni* (Fig. S3). The results showed that the dominant OPUs were closer to *Acinetobacter* sp. compared to the *Comamonas* sp., implying *that Acinetobacter* sp. are more likely the dominant cell in OPU1 and likely the potential functional microorganisms involved in PDN (Fig. S3). Moreover, the nitrification and denitrification rates in this study were also quantitively comparable with the pure culture study results of *Acinetobacter junii* (Ren et al., 2014), which had the highest similarity (>98%) to the representative OTU of *Acinetobacter* genus retrieved from our sample via sequence alignment. The ammonium oxidation rate in the nitrification batch activity tests was calculated as 2.39 ± 0.67 mg N/(g SS·h), which was in the range of 1.71 - 4.27 mg N/(g SS·h) of the *Acinetobacter junii*, and the nitrate reduction rate in the denitrification batch activity tests was 2.04 ± 0.47 mg N/(g SS·h), which was lower than the pure culture of 4.20 mg N/(g SS·h) probably explained by the lower C/N ratio (< 5) applied. Overall, we deduced that partial denitrification led to the nitrite accumulation in the B-stage and PHA played the role as the potential internal carbon source. Although further confirmation needed with more investigations. Nevertheless, these results, for the first time, provided direct and cellular level evidence that PHA maybe the potential internal carbon of the dominated bacteria clustered in OPU1 to support the partial denitrification process in absence of external carbon.

**Fig. 7.**
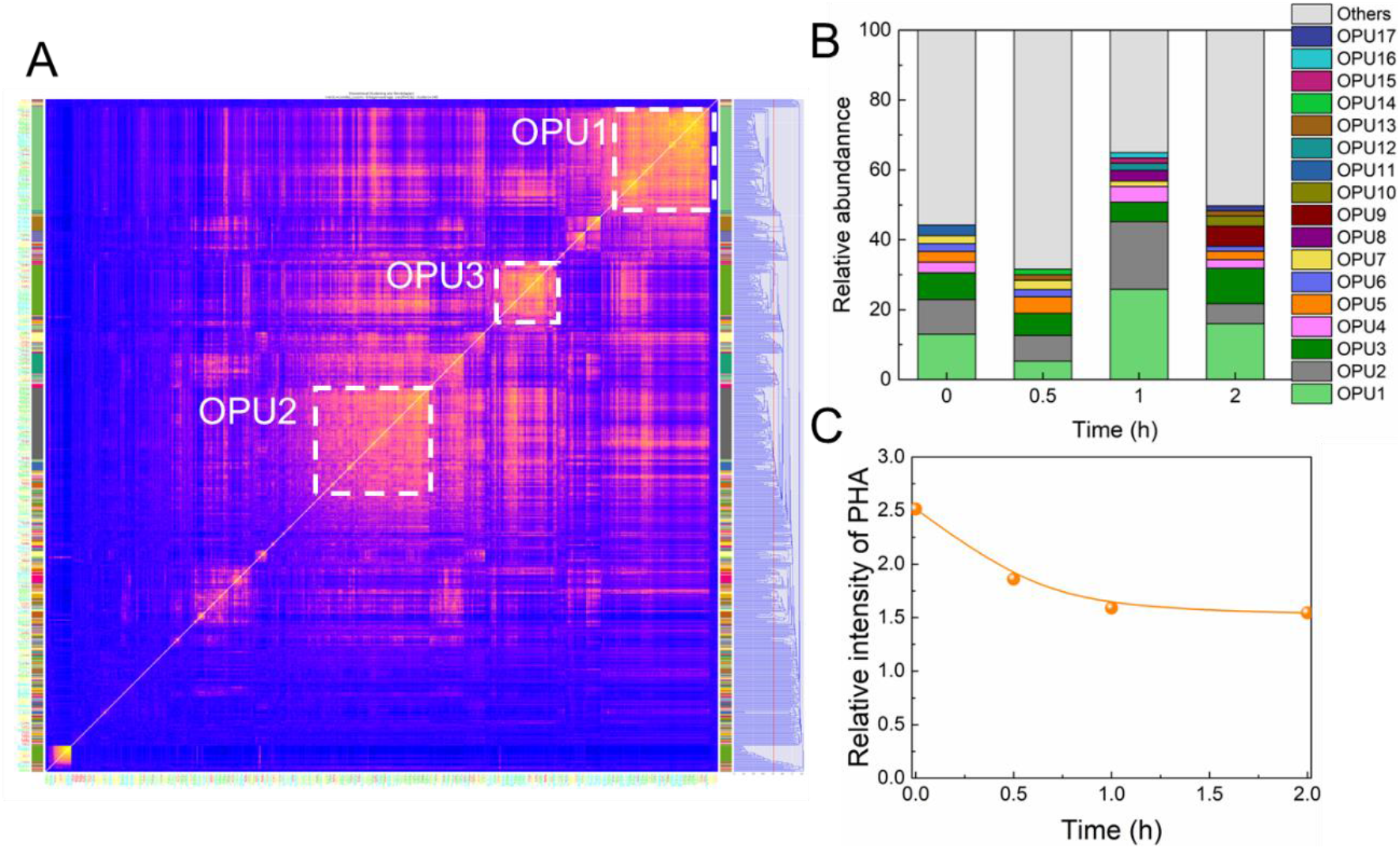
Single-cell Raman spectroscopy-based phenotyping analysis of the microbial communities during the denitrification batch activity test with fermentate pre-incubation. (A) HCA analysis of single-cell Raman spectra retrieved from different time points, using cosine similarity matrix and cut-off threshold of 0.82; (B) relative abundance of different clustered OPUs; (C) the changes in the relative intensity of PHA in the cells of dominant OPU1.

### 3.5. Significance and implications of this study

In this study, the new concept of integrating sidestream EBPR (S2EBPR) to the sustainable mainstream A-B process was tested in a pilot scale and the dominant microorganisms were identified by the combination of phylogenetic and phenotypic analysis for the first time. To enable EBPR that otherwise would be considered infeasible with A-B, the fermentate produced from the A-stage sludge digestion allowed carbon redirection in a more controllable manner to the S2EBPR reactor. Mass balance and SCRS-based phenotypic analysis suggested that the innovative configuration of the A-B-S2EBPR system seemed to enrich for the internal-carbon-containing heterotrophic bacteria. Nitrite accumulation induced by internal carbon-driven partial denitrification was evidenced to be the major nitrogen transformation pathway, which was combined with the downstream anammox to achieve efficient complete nitrogen removal for municipal wastewater. Unlike the previous method to achieve PDN via limiting external carbon to nitrogen ratio, which can often be infeasible to emply in practice, our study demonstrated that the excess external carbon via sludge fermentation could be transformed to the internal carbon source (i.e., PHA) in the S2EBPR reactor, and then served as the electron donor to drive the slow nitrate reduction over nitrite, which will facilitate the PDN process. This study could provide us a new strategy to utilize the sidestream sludge hydrolysis and fermentation to enrich the internal-carbon-containing heterotrophs to solve the long-existing SRT contradiction between autotrophic nitrifiers and heterotrophic PAOs, and finally achieve the simultaneous biological nitrogen and phosphorus removal.

Microbiologically, the exact functional organisms performing the partial denitrification have not been identified till now. Different from the known potential partial denitrifiers (*Thauera* and *Halomonas*) detected in the PDN systems, our pilot system enriched the genera of *Acinetobacter* (17.8 ± 15.5)% and *Comamonadaceae* (6.7 ± 3.4%), with the very low level of conventional nitrifiers detected (< 0.2%). The species *Acinetobacter junii* were previously reported to exhibit the ability of efficient N- and P-removal, respectively (Han et al., 2018; Ren et al., 2014). Moreover, rather than the common chemical analysis of the overall PHA variation, we applied the single-cell Raman spectroscopy technique to accurately reveal the existing dominant microorganism (the potential *Acinetobacter* sp.) in the A-B-S2EBPR system that contained the intracellular PHA, which could be the potential electron donor to drive the PDN. However, the phylogenetic identity and the functional ability to denitrify of the PHA-accumulating bacteria still needs to be investigated in depth via the joint analysis of the single-cell sorting based genomics.

## 4. Conclusions

This study, for the first time, demonstrated an efficient short-cut nitrogen removal via nitrite that was provided mostly from internal carbon-driven partial denitrification in a pilot A-B-S2EBPR system for treating municipal wastewater. The following conclusions were obtained:

1. Nitrite accumulation was detected in B-stage intermittently aerated tanks with the effluent concentration and NAR of 5.5 ± 0.3 mg N/L and 79.1 ± 6.5%, respectively. The effluent TIN concentration and removal efficiency of the A-B-S2EBPR system could reach 4.6 ± 1.8 mg N/L and 84.9 ± 5.6%.
2. Nitrite accumulation was proved to be caused mostly by partial denitrification process without NOB out-selection as confirmed by both activities assessment and microbial analysis of AOB and NOB.
3. The integration of S2EBPR could largely reshape the nitrogen microbial communities and enrich the internal carbon-accumulating candidate denitrifying genera including *Acinetobacter* (17.8 ± 15.5%) and *Comamonadaceae* (6.7 ± 3.4%).
4. PHA was approved to be the potential internal carbon source for partial denitrification as evidenced by phenotypic analysis and intracellular PHA analysis at single-cell and OUP-level via Raman spectroscopy.

## Supplementary Information

E-supplementary data of this work can be found in online version of the paper.

## Acknowledgements

This study was funded by the Water Environment Research Foundation (WRF 4901). D.K. was a visiting scholar supported by the fellowship of China National Postdoctoral Program for Innovative Talents (BX2021023) and China Scholarship Council (CSC). Special thanks are given to the generous support by operators and staff of Hampton Roads Sanitation District (HRSD). We also want to thank all the team members for the WRF project including Jim McQuarrie, Issac Avila, Dan Freedman, Rudy Maltos (Denver Metro Water Recovery), Beverley M. Stinson (AECOM), Christine deBarbadillo (DC Water), Paul Dombrowski (Woodard & Curran), Daniel Dair, Chandler Johnson (World Water Works Inc.) and James Barnard (Black & Veatch).

