## Supplementary information for "Molecular Evidence of Internal Carbon-Driven Partial Denitrification Annamox (PdNA) in a mainstream Pilot A-B System Coupled with Side-stream EBPR treating municipal wastewater"

Figures: 3

Tables: 2

Texts: 1

Pages: 10

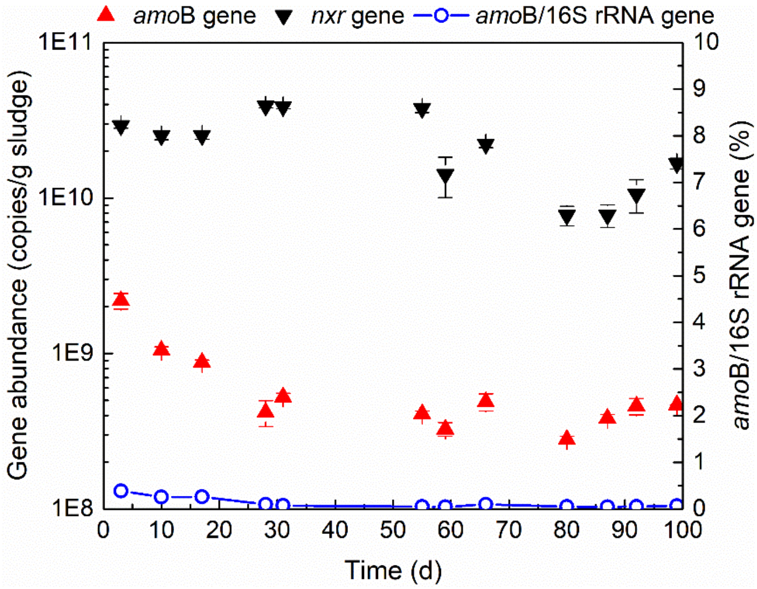


**Fig. S1** Gene abundance of Comammox (*amo*B gene) and NOB (*nxr* gene).

The average *amo*B gene abundance of Comammox was (6.55±5.31)×$10^{8}$ gene copies/g sludge, which was two magnitudes lower than the *nxr* gene abundance of *Nitrospira* as (2.29±1.17)×$10^{10}$ gene copies/g sludge. Moreover, the relative abundance of Comammox calculated as the *amo*B to 16S rRNA gene ratio was only 0.12±0.11%.

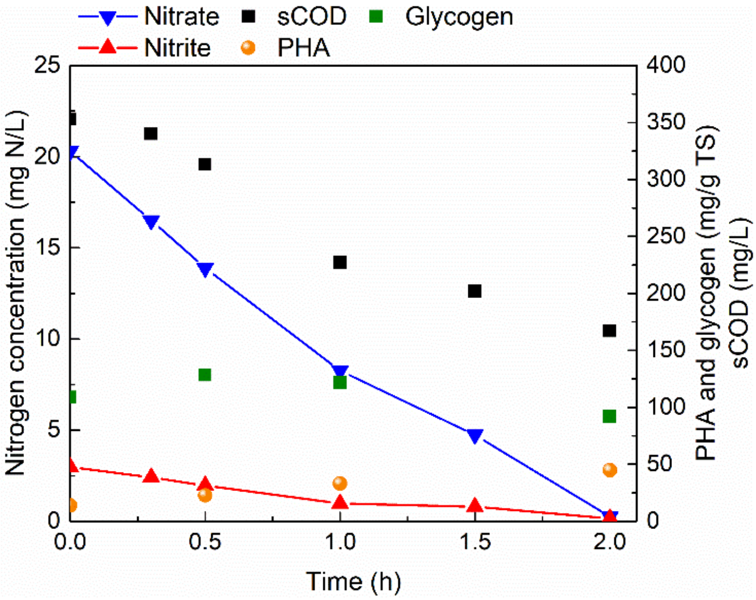


**Fig. S2** Denitrification batch activity test with acetate added as the external carbon source.

The initial acetate concentration was 350 mg COD/L with COD/$NO_3^-$ ratio as 17 and no nitrite accumulated during the whole process, and PHA accumulated gradually from 13.90 mg/g TS to 44.78 mg/g TS.

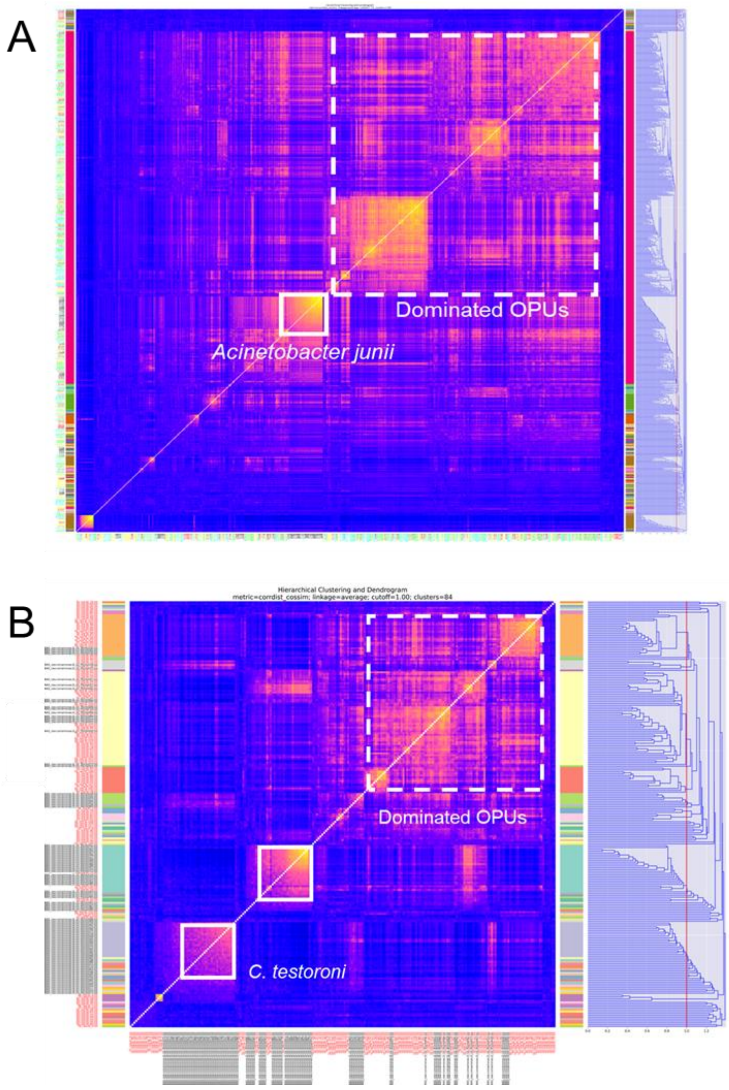


**Fig. S3** Hierarchical clustering analysis of the Raman spectra of *Acinetobacter junii* (A) and *Comamonas testosteroni* (B) with those retrieved from the sludge samples in fermentate pre-incubated denitrification batch activity test.

The single-cell Raman spectroscopy (SCRS) spectra of *Acinetobacter junii* and *Comamonas testosteroni*, which were identified as the species of *Acinetobacter* and *Comamonadaceae* that had highest similarity (>98%) to those sequences retrieved from our pilot system, were clustered with the dominate OPUs in our samples.

**Table S1** Operational and performance parameters of the pilot-plant.

| | **A-stage effluent** | **A-stage WAS fermentate** | **SBPR** | **B-stage** | | | | | | **MBBR** |
|---|---|---|---|---|---|---|---|---|---|---|
| | | | | **AN** | **AE1** | **AE2** | **AE3** | **AE4** | **Effluent** | |
| **Volume** (L) | - | - | 174 | 53 | 150 | 150 | 150 | 150 | - | 340 |
| **Flow rate** (L/min) | 2.4 ± 0.1 | 0.1 | 0.8 ± 0.1 | | | 5.3 ± 0.4 | | | 2.5 ± 0.1 | 2.5 ± 0.1 |
| **HRT** (h) | - | - | 3.7 | | | 5.1 ± 0.4 | | | - | 5.8 ± 0.2 |
| **SRT** (d) | - | - | - | | | 8.3 ± 1.3 | | | - | - |
| **RAS return and split ratio** (%) | - | - | 25 | | | 121.6±15.5 | | | - | - |
| **Aerobic fraction** | - | - | - | - | | 0.6 ± 0.1 | | | - | - |
| **pH** | - | - | 6.6 ± 0.3 | 7.0 ± 0.2 | | 7.1 ± 0.1 | | | - | 7.0 ± 0.2 |
| **Temperature** | - | - | 20.8 ± 1.4 | 20.8 ± 1.0 | | 20.9 ± 1.0 | | | - | 21.0 ± 1.2 |

| | | | | | | | | | | |
|---|---|---|---|---|---|---|---|---|---|---|
| (℃) | | | | | | | | | | |
| **TSS** (g/L) | - | - | 11.9 ± 1.6 | | | 4.3 ± 0.7 | | | 22.7 ± 10.3 | 21.2 ± 7.3 |
| **tCOD** (mg/L) | 222.7 ± 61.2 | 3923.2 ± 1178.6 | - | | | - | | | 64.8 ± 17.3 | 56.8 ± 10.7 |
| **sCOD** (mg/L) | 98.3 ± 28.5 | 1800.9 ± 404.0 | 93.6 ± 34.1 | 57.5 ± 11.6 | | | - | | 39.3 ± 6.1 | 35.0 ± 6.0 |
| **VFAs** (mgCOD/L) | - | 691.6 ± 167.3 | 30.5 ± 9.4 | | | - | | | - | - |
| **$NH_4^+$** (mg N/L) | 30.9 ± 3.7 | 116.2 ± 38.0 | - | 19.0 ± 3.0 | 15.5 ± 1.9 | 12.0 ± 2.0 | 8.6 ± 2.2 | 6.0 ± 1.5 | 6.5 ± 1.4 | 2.4 ± 1.4 |
| **$NO_2^-$** (mg N/L) | - | - | - | 0.7 ± 1.3 | 1.2 ± 0.6 | 2.2 ± 0.7 | 3.2 ± 1.2 | 3.7 ± 1.5 | 3.5 ± 1.3 | 0.6 ± 0.3 |
| **$NO_3^-$** (mg N/L) | - | - | - | 0.4 ± 0.2 | 1.2 ± 1.0 | 2.6 ± 1.7 | 4.0 ± 2.5 | 5.1 ± 2.3 | 5.0 ± 2.9 | 1.5 ± 1.2 |

**Table S2** Sequence and thermocycling conditions for Primer sets of qPCR

| | Primer | Sequence (5'-3') | Condition | Reference |
|---|---|---|---|---|
| Total bacteria | 341F/534R | CCTACGGGAGGCAGCAG/ATTACCGCGGCTGCTGGCA | 95°C for 3 minutes, then 40 cycles of 94°C for 30 seconds, 60°C for 45 seconds, and 72°C for 30 s. | (He et al., 2007) |
| AOB | amoA-1F/2R | GGGGTTTCTACTGGTGGT/CCCCTCKGSAAAGCCTTCTTC | 95°C for 3 minutes, then 40 cycles of 95°C for 15 seconds, 54°C for 30 seconds, and 72°C for 1 minutes. | (Rotthauwe et al., 1997) |
| *Nitrospira* | nxrB-169F/638R | TACATGTGGTGGAACA/CGGTTCTGGTCRATCA | 95°C for 5 minutes, then 35 cycles of 95°C for 40 seconds, 56.2°C for 40 seconds, and 72°C for 90 seconds. | (Pester et al., 2014) |
| Comammox | amoB-148F/485R | TGGTAYGAYACNGAATGGG/CCCGTGATRTCCATCCA | 95°C for 3 minutes, then 40 cycles of 95°C for 15 seconds, 52°C for 45 seconds, and 72°C for 1 minutes. | (Cotto et al., 2020) |

**Text S1** Raman spectra data preprocessing methods

Load the data into software LabSpec 6. Split the array to get the spectrum of each sampling points. Combine the data of ~10 background points with adjusting overlapping intensities to produce the average background spectrum. Screen the data for removing cosmic ray/spike, outlier and poor-quality spectra. Conduct a smoothing, filtering and correction algorithm to filter high-frequency noise in spectra (Size 3, Degree 4, Polynomial type, Correction 1). Conduct a baseline correction algorithm to account for fluorescence interference and subtract averaged background from sample spectrum (Polynomial type, Degree 10, Max points 256, Noise points 64). Find the peaks of the processing spectrum (Amplitude 5%, Size 20 pix). Identify the single-cell spectrum by removing the spectrum with the intensity of either 995-1010 $cm^{-1}$ (bacteria's phenylalanine) or 1640-1690 $cm^{-1}$ (bacteria's amide I) as 0.
